# Augmented Reality and Cross-Device Interaction for Seamless Integration of Physical and Digital Scientific Papers

**DOI:** 10.1101/2024.02.05.578116

**Authors:** Md Ochiuddin Miah, Jun Kong

**Author notes:** Corresponding author *Email address:* (Md Ochiuddin Miah).

## Abstract

Researchers face the challenge of efficiently navigating vast scientific literature while valuing printed papers in the digital age. Printed materials facilitate deeper engagement and comprehension, leading to superior exam performance and enhanced retention. However, existing digital tools often need to pay more attention to the needs of researchers who value the tactile benefits of printed documents. In response to this gap, we introduce AR-PaperSync, a transformative solution that leverages Augmented Reality (AR) and cross-device interaction technology. AR-PaperSync seamlessly integrates the physical experience of printed papers with the interactive capabilities of digital tools. Researchers can effortlessly navigate inline citations, manage saved references, and synchronize reading notes across mobile, desktop, and printed paper formats. Our user-centric approach, informed by in-depth interviews with six researchers, ensures that AR-PaperSync is tailored to its target users’ needs. A comprehensive user study involving 28 participants evaluated AR-PaperSync’s significantly improved efficiency, accuracy, and cognitive load in academic reading tasks compared to conventional methods. These findings suggest that AR-PaperSync enhances the reading experience of printed scientific papers and provides a seamless integration of physical and digital reading environments for researchers.

## 1. Introduction

A vast expanse of scientific literature increasingly dominates the landscape of academic research. This growth necessitates an efficient modality for researchers to assimilate scholarly content. Traditionally, the academic community has shown a strong inclination towards printed scientific papers, attributed to their tangible nature facilitating deeper engagement and comprehension [1, 2]. This preference is substantiated by empirical studies indicating superior exam performance, enhanced retention, and active reading engagement when using printed materials compared to digital formats [3, 1, 2]. However, the advent of digitalization in academic research presents a dichotomy—the tactile benefits of print media versus the dynamic and interactive capabilities of digital tools [2].

Current systems predominantly focus on digital enhancements [4, 5, 6] or AR-driven immersive reading experiences presenting visual content [7, 8, 9], often neglecting the unique requirements of researchers who value the physicality of printed documents.

Despite advancements in digital reading technologies, there remains a significant gap in solutions that effectively integrate the physical experience of printed papers with the convenience and interactivity of digital tools. While adept at enhancing digital reading experiences [10, 11, 12], current research systems need to address the challenges faced by researchers relying on printed materials. These challenges include efficiently navigating inline citations, managing saved citations, and synchronizing reading notes across various formats [4, 5, 13]. Researchers often face the daunting task of manually tracking and cross-referencing inline citations. This process can be time-consuming and disruptive to the natural flow of reading and comprehension. Also, managing saved citations in printed formats requires a lot of work. Traditional methods usually involve manual cataloguing and organization and need more efficiency and flexibility than digital tools offer [5, 6]. Such methods are often labor-intensive and can lead to fragmented research processes, hindering the researcher’s productivity and effectiveness. Another significant challenge lies in synchronizing reading notes and insights across printed and digital formats [13]. Researchers are often required to maintain separate systems for note-taking and reference management for their printed and digital papers. This leads to disjointed workflows, increased cognitive load, and a higher likelihood of critical insights being overlooked or lost in the transition between formats.

Our motivation stems from the need to cater to researchers who prefer printed papers while recognizing the benefits of digital enhancements. Studies have shown that reading from printed papers leads to higher exam performance [3], comprehension, and retention compared to digital mediums [1, 2]. Therefore, we focus on developing a solution that respects and enhances traditional reading practices, bridging the gap between the physical and digital realms. To address researchers’ challenges while reading printed papers, we introduce AR-PaperSync, an innovative solution for mobile and desktop that leverages Augmented Reality (AR) and cross-device interaction technology (Figure 1). AR-PaperSync represents a pivotal step towards revolutionizing how researchers interact with scholarly content. By seamlessly integrating digital enhancements into the physical realm of printed papers, the AR-PaperSync mobile application offers a transformative reading experience that caters to traditionalists’ preferences while harnessing modern technology’s capabilities.

**Figure 1:**
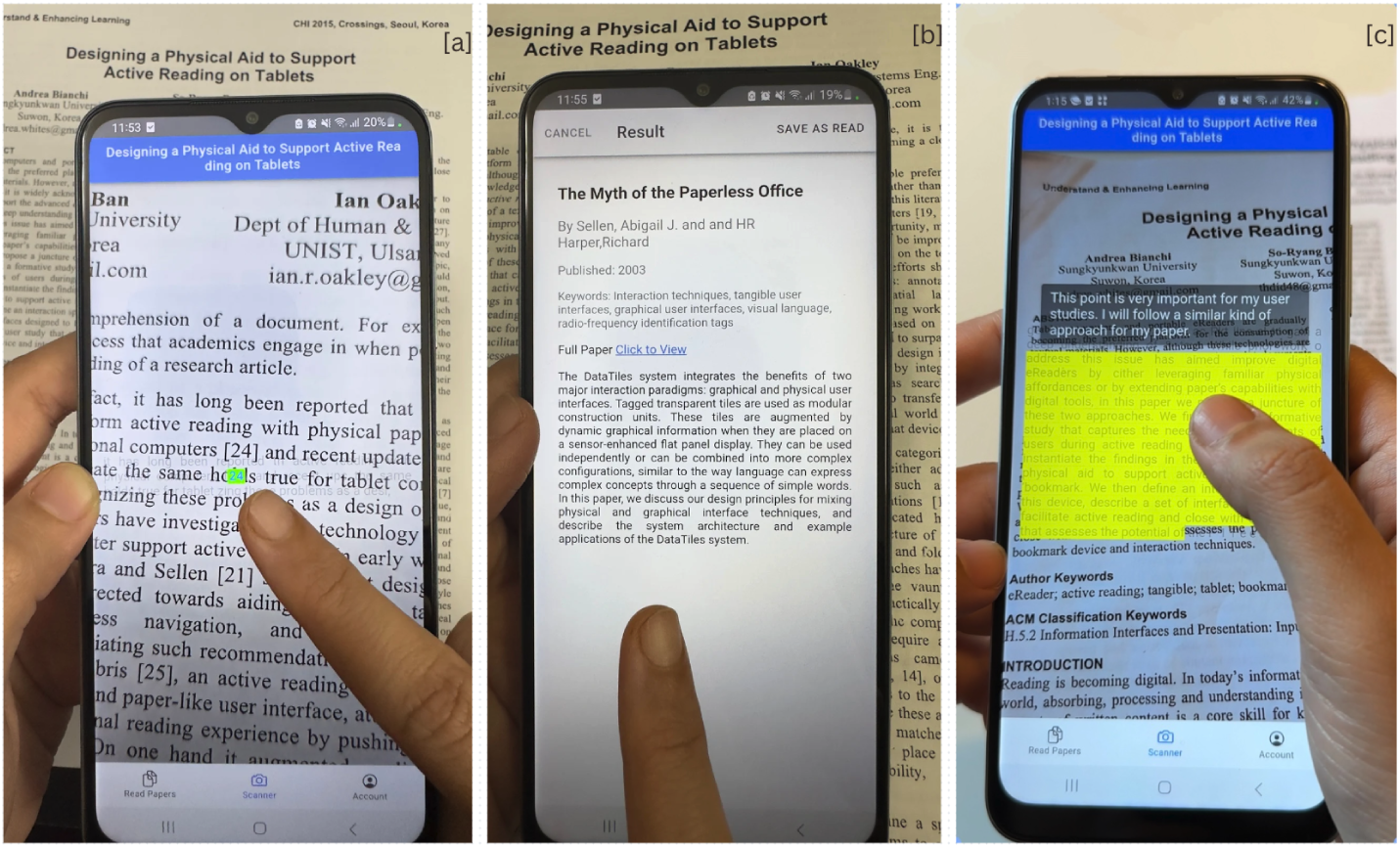
Overview of AR-PaperSync. (a) enabling users to access citations from physical documents through a mobile AR application effortlessly; (b) viewing full paper and tracking saved inline citations on physical paper; (c) enabling real-time synchronization across mobile, desktop, and printed paper formats, ensuring continuity in their research activities.

At the core of AR-PaperSync lies a commitment to enhancing the reading journey for researchers who treasure the tangible qualities of printed papers. Inline citations, often frustrating and interrupting the reading process, become effortless to navigate with AR-PaperSync. Through AR overlays, researchers can instantly access additional information about citations, view related references, and explore cited works in a fluid and non-disruptive manner. Once laboriously cataloged, saved citations are managed seamlessly within the AR-PaperSync ecosystem. The system intelligently recognizes and organizes saved references, enabling researchers to access detailed bibliographic information with a simple gesture. Furthermore, AR-PaperSync addresses the challenge of reading note synchronization by providing a unified platform for researchers to create, manage, and sync their notes across mobile, desktop, and printed paper formats. This synchronization enhances the researcher’s productivity and ensures that valuable insights and annotations are preserved, irrespective of the format.

Our development of AR-PaperSync is guided by a user-centric approach, informed by in-depth interviews conducted with 6 researchers representing diverse academic backgrounds. This interview ensures the tool is meticulously tailored to its target users’ needs and operational processes. Subsequently, we conducted a comprehensive user study involving 28 participants to assess AR-PaperSync’s practical utility in real-world scenarios. The evaluation encompassed the tool’s effectiveness in facilitating seamless navigation of inline citations, efficient management of saved references, and the seamless sharing of reading notes across various formats. The outcomes of the study indicate that AR-PaperSync significantly improves the efficiency and accuracy of navigating citations and synchronizing notes across mobile, desktop, and printed paper formats. Analysis of usability questionnaires, including the System Usability Scale (SUS) and the NASA Task Load Index (NASA-TLX)), underscores AR-PaperSync’s remarkable capacity to reduce cognitive burdens and ease of integration into researchers’ workflows.

In explicit terms, this paper presents the following key contributions:

- Introduce an innovative scientific paper inline citation searching tool—AR-PaperSync—that overcomes existing limitations by seamlessly integrating physical and digital versions of scientific papers using Augmented Reality (AR) and cross-device interaction. Unlike prior approaches focusing on either reading articles [14, 6, 11, 12] or navigating citations [4, 5] on digital platforms, AR-PaperSync harmonizes the two, enabling users to access citations from physical papers through a mobile application effortlessly.
- AR-PaperSync enables real-time synchronization across mobile, desktop, and printed paper formats, ensuring continuity in their research activities. By facilitating syncing reading notes from digital to physical paper and tracking saved inline citations on physical paper, AR-PaperSync provides a smooth and consistent user reading experience.
- Insights gathered from interviews with (N=6) researchers formed the basis for AR-PaperSync’s user-centric design to ensure the tool is meticulously tailored to its target users’ needs and operational processes.
- Validation of AR-PaperSync’s effectiveness through a user study involving (N=28) participants demonstrated its proficiency in efficiently navigating and tracking inline citations from printed papers and seamlessly connecting various reading environments to share reading notes.

## 2. Related Work

Previous research has focused on improving the digital reading experience of scholarly content. Several systems have been developed for digital readers to navigate referenced scientific documents [4, 15, 16], exploring relevant prior work [5, 17, 18], linking sentences with table cells to enhance interactivity [6], and showing definitions of technical terms and symbols [11]. Additionally, AR technologies have been employed to create immersive reading experiences [7], including presenting paragraphs, scientific data and visual content [9, 8]. However, existing works still need a specific focus on seamlessly integrating physical and digital versions of scientific papers, a critical aspect our proposed tool, AR-PaperSync, aims to address.

### 2.1. Systems for Reading in Digital Space

Numerous systems have been developed to enhance the reading experience of scientific papers in digital formats [19, 16, 10]. These systems often improve text readability, provide contextual information, explore prior work and content navigation. For instance, Park et al. [4] developed QuickRef, an interactive reader, to assist scholars in navigating referenced documents that provide additional information about cited papers. It addresses the challenges of navigating cited papers by presenting meta-information, an overview, and relevant information. Additionally, Chang et al. [5] introduced a personalized literature review tool, CiteSee, to enhance the reading experience of scientific papers. It leverages a user’s previous research activities to provide contextual information and facilitate exploring relevant prior work. Kim et al. [6] conducted a user study comparing their interactive document reader, which links sentences to table cells, with a baseline reader. This study highlights the potential of interactive document readers to reduce split attention and improve document reading. Additionally, Head et al. [11, 12] introduced an augmented reading interface, pioneering innovations like tooltips for position-sensitive definitions, decluttering filters, and novel visual design patterns for mathematical notation, aiming to improve the readability and comprehensibility of research papers in digital formats. However, it is worth noting that while these endeavours have predominantly focused on enhancing the digital reading experience, there is evidence suggesting a preference for printed articles over e-books among readers [3].

In light of this, our research takes a different approach by prioritizing enhancing the reading experience for printed paper readers. We achieve this by developing a smartphone-based AR system that supports navigation and facilitates tracking of inline citations while providing digital reading notes on printed documents, offering a novel solution for improving the traditional reading experience.

### 2.2. Reading Scientific Papers using Augmented Reality

Augmented reality (AR) has transformed many areas of research and education. Some AR systems are designed to visualize scientific data, simulations, images, and user interfaces [20, 9, 7, 21]. Rzayev et al. [7] studied how different text presentations (like RSVP and paragraph) and text placements (world-fixed, edge-fixed, head-fixed) impact the VR environment. Meanwhile, Al-Ali et al. [9] introduced an augmented interactive reality (AIR) tool named ”My Vision-AIR.” It’s designed to enrich adult reading experiences by infusing AR into traditional books. This app uses 2D/3D graphics, animations, virtual buttons, translations, and multimedia to make reading more engaging and immersive. On another note, Bianchi et al. [14] launched eTab, a tool to elevate the tablet reading experience. Kirner et al. [20] contributed to the AR landscape by developing the interactive book GeoAR, leveraging Augmented Reality for teaching Geometry topics. Meanwhile, Rajaram et al. [22] introduced Paper-Trail, a transformative tool designed to enrich paper-based learning with digital media, animations, and clipping masks. These AR applications have significantly advanced text presentation, enhanced the reading experience for adult learners, facilitated image visualization, and improved the teaching of geometric shapes. However, our research focuses on a unique aspect within the AR domain, addressing the needs of researchers by facilitating the navigation of inline citations, tracking saved references in printed papers, and seamlessly updating reading notes between physical and digital environments. Our work bridges the gap between traditional and digital reading experiences by ensuring a consistent reading journey, offering valuable contributions to the field.

### 2.3. Integrating Scientific Papers with Digital Environments

As technology evolves, a growing interest has been syncing traditional paper documents with digital environments. Efforts have been made to bridge the gap between traditional printed content with digital enhancements—mobile, desktop—facilitating real-time synchronization and interaction across platforms [23, 24, 25, 26]. Among these, Qian et al. [13] developed an AR-based system to enhance interactions with printed documents through layout recognition, leading to more accurate and efficient annotations. Steimle et al. [27] created CoScribe, which integrates pen-based annotations on physical documents with digital versions. Similarly, Guimbretìere [28] introduced PADD (Paper Augmented Digital Documents), blending paper and digital interactions via patterns recognized by a digital pen. Motoki and Koike [29] also contributed with ”EnhancedDesk,” a desktop interface merging paper with digital information through 2D matrix codes. These developments highlight the potential of combining tangible and digital media using various tools like pens, tabletops, or projectors. Building on this foundation, we have developed AR-PaperSync, an AR mobile application designed to enhance the reading experience, especially for scholars accustomed to traditional scientific papers. This innovative app reduces the need for additional devices and uses AR to connect printed documents with their digital counterparts fluidly. It simplifies access to inline citations, tracks saved references on physical papers, and updates reading notes across mobile, desktop, and printed formats through AR and cross-device interaction.

## 3. Interview

In the initial phase of the AR-PaperSync project, we conducted interviews to gain insights into how researchers interact with inline citations and reading notes during their academic reading and to identify common limitations. Understanding these aspects was crucial for developing targeted design goals for AR-PaperSync. We recruited 6 participants with diverse educational backgrounds, including 3 PhD candidates, 2 Master’s, and 1 Bachelor’s student from transportation and logistics, microbiology, computer science, biochemistry, and mechanical engineering. The interviews, lasting approximately 20 minutes each, followed a semi-structured format. We focused on three main questions to uncover participants’ reading habits, technology usage, and challenges faced while interacting with printed scientific papers, with follow-up questions posed to clarify their responses.

### 3.1. Reading Preferences and Practices

The participants’ responses revealed various reading preferences, emphasizing the varied reliance on printed and digital papers. One participant highlighted a strong preference for the *”tangibility of printed papers”* for in-depth research, often engaging with *”3-6 papers per week”*. This preference for physical copies was echoed by another who *”prefers printed papers for initial reading”* and consistently reads *”3-4 papers weekly in printed form”*. In contrast, some participants balanced their use of digital and printed formats, with one noting the practice of printing *”complex papers for better comprehension”* while generally favoring digital formats for convenience. Another described printing papers primarily for *”group discussions”*, indicating a situational reliance on physical documents. These insights underscore the diverse academic reading habits, ranging from a predominant preference for printed materials for detailed study to a strategic combination of digital and physical mediums.

### 3.2. Use of Technology with Printed Papers

The participants’ approaches to integrating technology with printed papers varied significantly, reflecting a broad spectrum of practices in academic research. Several participants described using a laptop as a complementary tool for their reading process. One participant specifically mentioned using a laptop for *”cross-referencing information and deeper analysis”*, highlighting the role of digital tools in enhancing the understanding of printed materials. Another researcher noted the use of a laptop for *”searching and saving relevant citations on printed paper”*, illustrating how digital tools facilitate the management of academic resources. In contrast, one participant pointed out that they *”rarely use a laptop in conjunction with printed papers”*, suggesting a preference for more traditional, paper-centric approaches. Interestingly, a dual-screen setup was mentioned for multitasking purposes, indicating that some researchers optimize their digital environment to augment their interaction with printed papers. A tablet was also noted for additional research, particularly when not at a desk, indicating a preference for more portable digital solutions. These varied responses underscore how researchers combine digital and physical resources to enhance these multifaceted reading and research habits. They also conclude it consumes time to integrate technology with printed papers during their research activities.

### 3.3. Managing Citations and Notes

The methods employed by participants for managing citations and notes while reading printed papers demonstrated a diverse range of practices. Several participants described manual strategies, such as checking inline citations by hand and using sticky notes for marking influential citations and thoughts. This traditional approach, while familiar, was often cited as time-consuming and sometimes led to challenges like misplacing physical papers. One participant specifically mentioned the frustration of losing papers, highlighting the limitations of purely manual systems. Others integrated digital tools into their workflow: one researcher used a digital tool for managing citations coupled with handwritten notes on paper margins, while another used a mobile app for scanning and organizing citations, occasionally complementing this with digital notes. Google Lens was also noted for checking citations, with saved information being explored later from a separate file. A mix of digital and manual methods was standard, with one participant typically using a desktop for note-taking at home but switching to manual checks and paper markings while on campus. These varied practices underline the need for a system that seamlessly integrates and enhances these methods, providing a more efficient and organized way to manage citations and notes.

### 3.4. Design Goals

The insights gleaned from the interviews with participants have directly informed the development of specific design goals for AR-PaperSync. These goals address the challenges and preferences in managing inline citations and notes and integrating digital tools with printed papers. The following design goals aim to enhance the academic reading experience by leveraging augmented reality (AR) and digital synchronization technologies:

- **[DG1]** Help researchers find inline citations from printed scientific papers connecting digital AR view using AR mobile applications. This goal aims to enhance the traditional method of accessing citations, improving overall reading efficiency, particularly for those who prefer printed materials.
- **[DG2]** Save inline citations on AR view for printed papers, offering a seamless way to manage and organize these references effectively.
- **[DG3]** Enables real-time synchronization of reading notes across printed paper, mobile, and desktop applications. This goal addresses the diverse use of technology and the challenges in managing citations and notes, aiming for a cohesive research process that integrates the physical and digital papers.

## 4. AR-PaperSync System

The development of AR-PaperSync represents a pioneering endeavor at the intersection of mobile augmented reality, digital document management, and real-time synchronization technologies. Here, we delve into the intricate architectural design and meticulous implementation that underpins the AR-PaperSync system, showcasing how it transforms the traditional reading experience into a dynamic and interactive journey.

### 4.1. Architectural Overview

The AR-PaperSync system is architecturally divided into three main components: the mobile application for AR clients, the desktop application for digital clients, and the back-end cloud service. Each component is meticulously designed to interact synergistically, providing a robust and user-centric experience. Figure 2 illustrates the architectural overview of AR-PaperSync.

**Figure 2:**
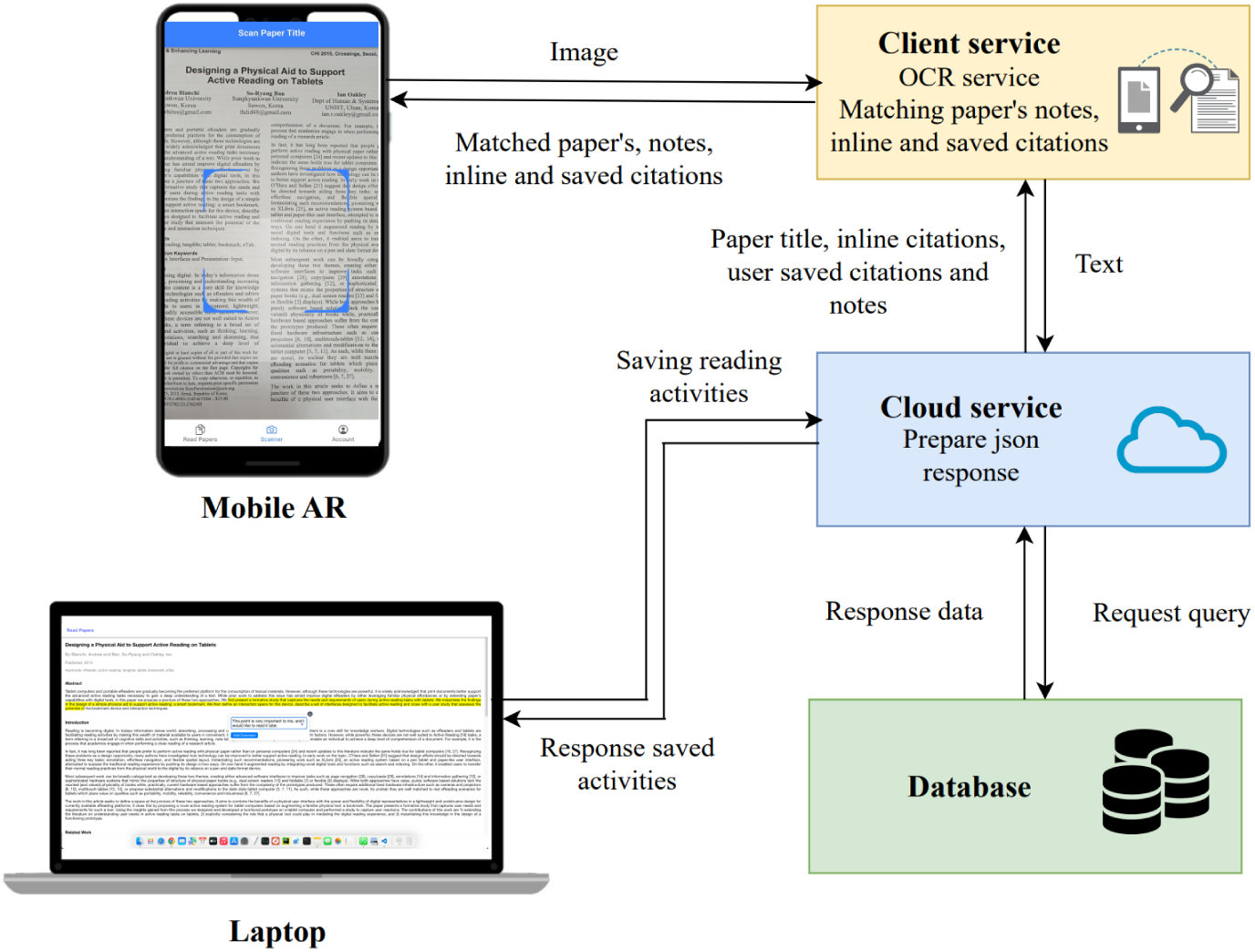
Architectural Overview. AR-PaperSync uses cloud and client service to facilitate the synchronization of matched paper titles, inline citations, reading notes, and saved citations across mobile applications (Mobile AR). Digital clients utilize desktop applications (Laptops) to create and store reading notes, with these notes being stored in the cloud database and synchronized in real time for mobile AR access.

### 4.2. Mobile Application for AR Client

The AR Client is meticulously developed using the Ionic 7 framework, Angular 16 for handling the application logic, and TypeScript 5 for scripting requirements. Capacitor 5 is a cross-platform bridge, enabling seamless access to native device features. The client service within the mobile AR application communicates over HTTP with the cloud service, utilizing JSON strings for efficient data exchange related to paper titles, inline citations, user-saved citations, and reading notes. Figure 3 encapsulates the core functionalities of the mobile AR, highlighting features such as the AR camera, paper title detection, inline citation display, and options to mark as read or access the entire paper.

**Figure 3:**
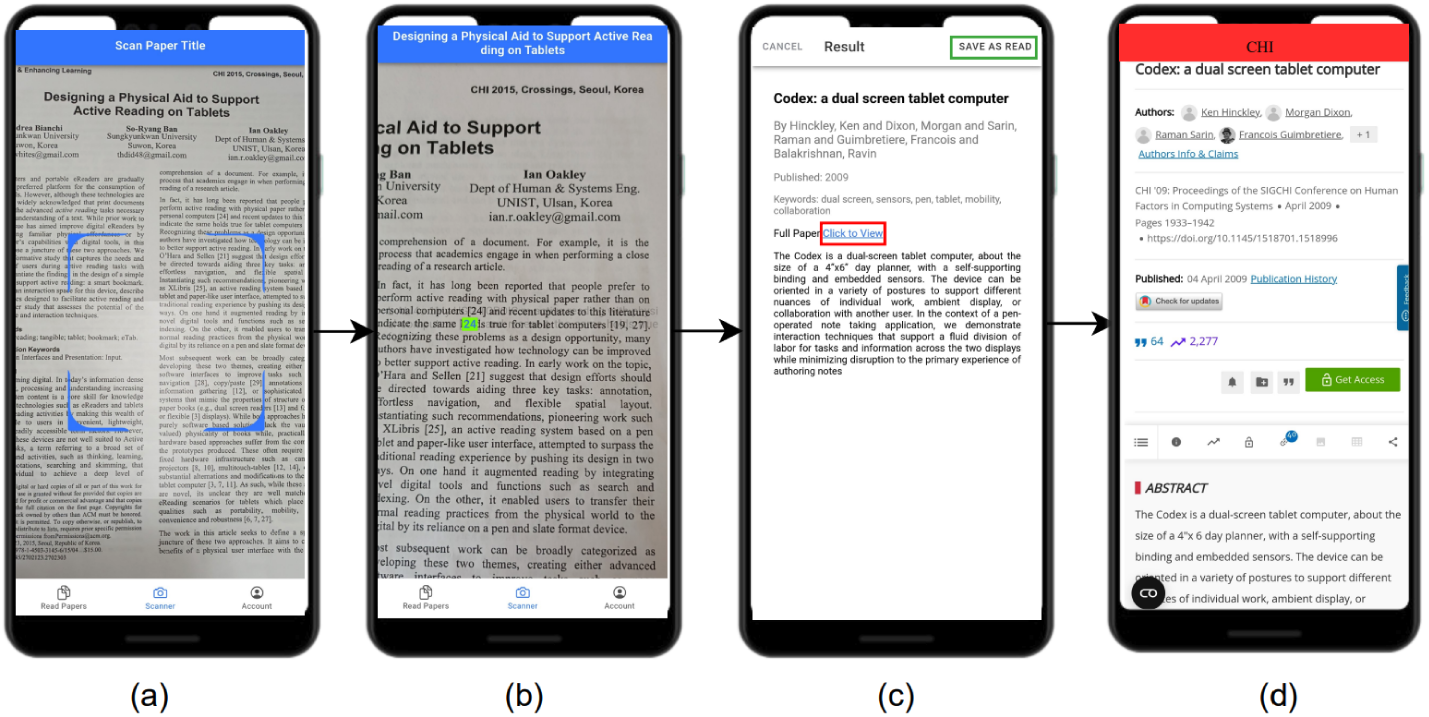
Mobile AR. (a) camera and AR module poised for text processing; (b) detection of paper title accompanied by inline citation, visualized within the AR view; (c) upon selecting augmented citations, detailed information becomes accessible, accompanied by options to mark as read or view the entire paper; (d) display of the complete paper sourced from the published site.

#### Camera and AR Module

The mobile application leverages the device’s camera and integrates augmented reality functionalities to detect and interact with physical papers effectively. This module initiates the document recognition process, captures images optimized for OCR processing, and overlays interactive AR elements on the physical paper. To achieve this, we utilized the capacitor camera preview to develop a customized camera interface compatible with Android and iOS platforms, incorporating AR views. Additionally, the ImageCompressService is implemented to compress captured images to a maximum of one megabyte, enhancing processing efficiency without compromising quality.

#### OCR Integration

Upon capturing the optimized image, the OCR module facilitates extracting and processing text content from the document, encompassing titles, inline citations, and reading notes. Given the criticality of accuracy and processing speed, a robust algorithm is imperative to effectively accommodate diverse fonts, sizes, and document conditions. To address this, we integrated Tesseract 4, leveraging its new neural net (LSTM) based OCR engine to extract text from images with unparalleled precision and efficiency.

#### User Interaction Layer

The user interaction layer is pivotal in rendering interactive elements on the user’s device, facilitating a compelling AR experience. Users can tap on augmented citations to access detailed information, navigate to cited papers seamlessly, or bookmark them for future reference. Moreover, this layer enables users to visualize reading comments associated with printed papers, enhancing their comprehension and engagement. Selected texts on printed papers are highlighted through the AR camera, and associated comments become accessible upon clicking the highlighted text within the AR view, fostering an immersive and interactive reading environment.

#### 4.2.1. Desktop Application for Digital Client

The Digital Client is an interactive platform that enables users to engage with digital renditions of academic papers while offering robust features for reading, annotating, and organizing documents. Developed utilizing Angular 16 and TypeScript 5, the Digital Client ensures a responsive and user-centric interface, facilitating seamless navigation and interaction. Figure 4 delineates the main components of the digital reading interface, emphasizing note-taking functionalities and synchronization capabilities with mobile AR.

**Figure 4:**
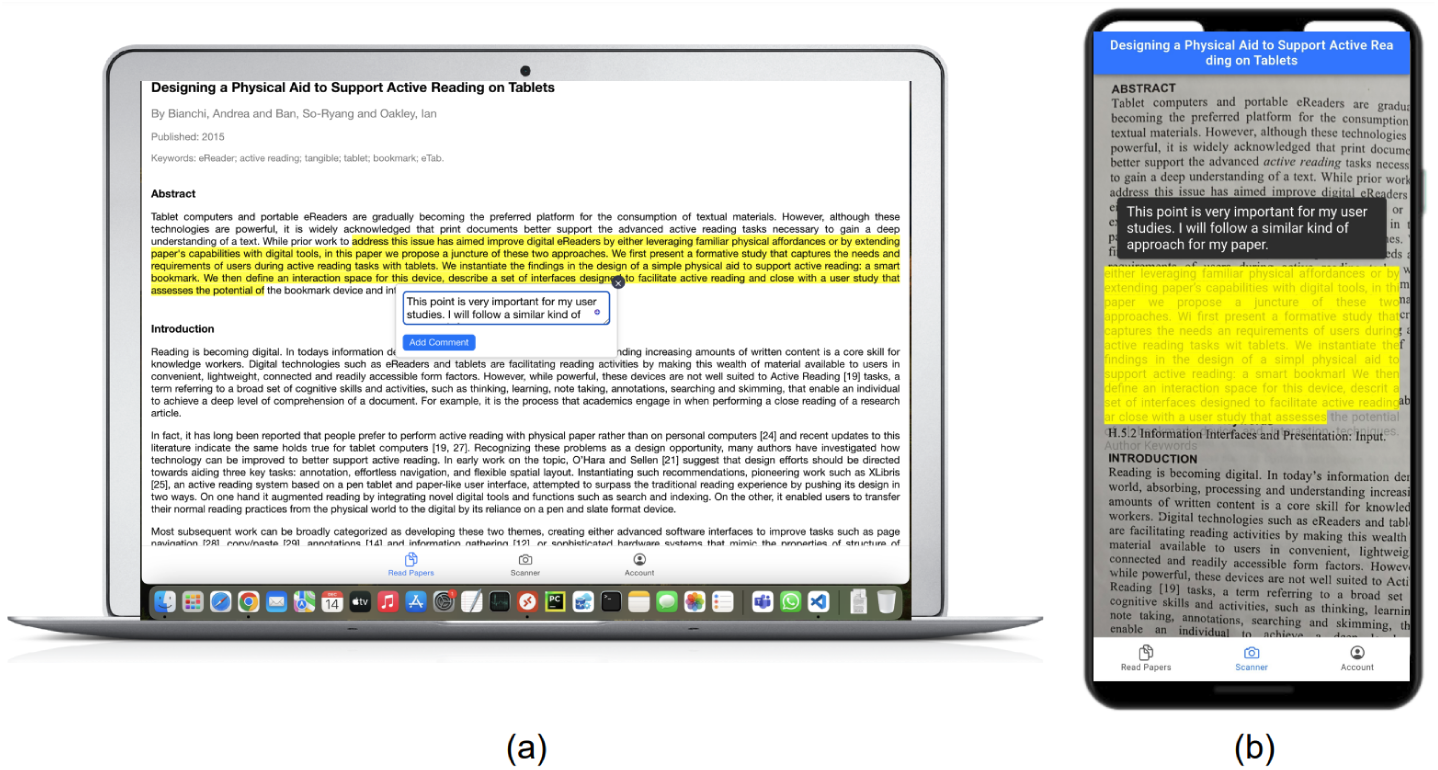
Digital Client Integrated with Mobile AR. (a) digital reading interface showcasing scientific papers with annotated notes, allowing users to add or modify annotations; (b) AR client enabling real-time access and interaction with annotations on printed papers, utilizing the AR camera’s capabilities.

#### Digital Reading Interface

The Digital Reading Interface is designed to present digital documents efficiently, accommodating various PDF formats while prioritizing an intuitive user experience. Leveraging libraries such as PDF.js enhances the presentation of PDF documents, ensuring clarity and accessibility. Users are empowered to annotate texts, add or modify reading notes, and organize content, thereby facilitating efficient academic engagement.

#### Note Synchronization with Mobile AR

The Digital Client facilitates a bidirectional data exchange with the Cloud Service Backend, ensuring synchronized interactions across digital and physical platforms. Notes, highlights, and anno-tations created within the Digital Client are seamlessly integrated with the AR Client, enabling real-time updates and reflections. The Digital Client communicates with the cloud service via HTTP, optimizing data exchange through JSON strings to synchronize users’ reading notes efficiently. Consequently, users engaging with the AR Client can access and interact with notes on printed papers in real time, leveraging the capabilities of the AR camera.

#### 4.2.2. Back-end Cloud Service

The Cloud Service Backend is a crucial element within the AR-PaperSync system, functioning as a centralized repository and a processing hub for user data, document information, and synchronization directives.

#### Database and Storage

Utilizing MS SQL Server, the backend efficiently manages a vast dataset encompassing user profiles, document metadata, annotations, and citations. The database design focuses on scalability, query efficiency, and data integrity. Specifically, we maintain essential paper details with associated citations in a structured manner. A dedicated citation table also establishes one-to-many relationships, ensuring comprehensive data organization. User-generated reading notes are systematically stored in a notes table, associating each entry with specific user and paper identifiers.

#### API and Data Services

The backend facilitates seamless interactions through RESTful APIs, leveraging Node.js to facilitate data exchanges between mobile and desktop clients. These services proficiently manage tasks such as storing citations, retrieving detailed paper and citation information, and synchronizing reading activities across platforms. Prioritizing user privacy and security, the backend incorporates robust measures, including authentication protocols and data encryption, safeguarding sensitive user information.

### 4.3. System Features

In realizing the objectives set forth for AR-PaperSync, we’ve successfully implemented three pivotal features: the exploration of inline citations, the saving of inline citations within the AR environment, and the presentation of reading notes on physical papers via augmented reality.

#### 4.3.1. [DG1] Explore Inline Citations using AR

Our primary design goal is to empower users to explore inline citations while reading printed papers. To retrieve citation details, readers simply scan the paper title using the AR camera view, obtaining real-time information. This establishes the foundation for acquiring citations from a specific paper. By scanning the content through the camera, users seamlessly access digital citations overlaid on the printed paper through our mobile AR application. Clicking on citations reveals detailed paper information, enhancing the reading experience with immediate access to supplementary materials and contextual insights. Figure 3 illustrates the step-by-step process of obtaining the paper title and its citations.

To optimize the citation exploration performance, real-time image compression occurs before sending the image to the OCR engine. The OCR service then converts the compressed image into plain text. If the application lacks a predefined paper title, it initiates a matching process with the paper storage. Upon successfully identifying the paper title, the application proceeds to search for citations within the text. Extracted citations are presented as clickable buttons on the AR view, accurately positioned based on the layout of printed paper citations. Readers can effortlessly click on these digital citations to access and explore detailed paper information. Figure 5 provides a visual representation of the workflow, illustrating how images are processed and seamlessly integrated into the AR mobile view.

**Figure 5:**
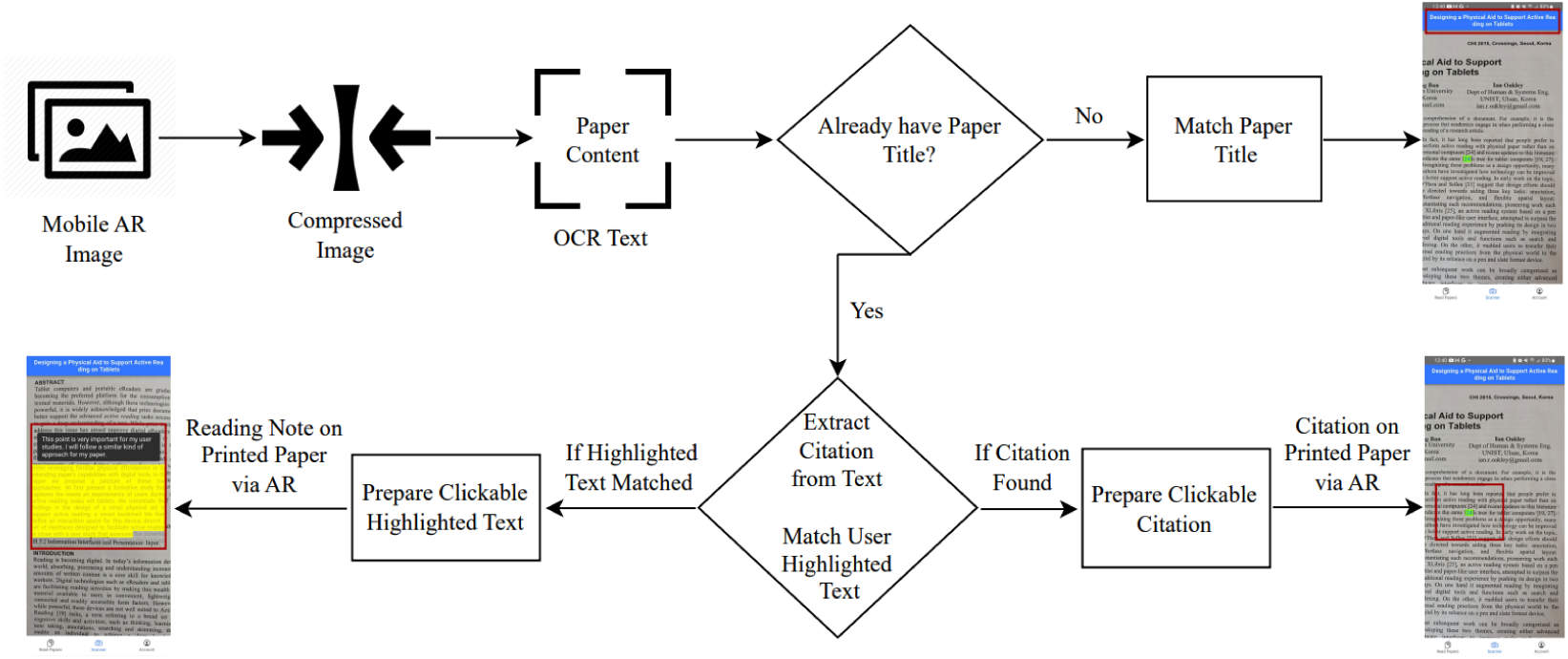
Detailing the exact sequence of image processing steps and their seamless integration into the AR mobile view, the system adeptly presents paper titles, inline citations, and reading notes.

#### 4.3.2. [DG2] Save Inline Citations on AR View

Building upon the foundation laid by exploring inline citations using AR, the AR-PaperSync system seamlessly integrates a feature allowing users to save identified inline citations for future reference. Upon successfully obtaining citation details through the AR camera view, users are presented with an intuitive ”Save as Read” button. Clicking on this button initiates the citation storage in the user’s reading activity database, enabling them to revisit and explore saved citations at their convenience.

The saved inline citations are stored securely within the cloud service backend, ensuring accessibility from any device and platform. As users continue scanning for inline citations, the AR view intelligently distinguishes between citations saved as read and those yet to be explored. Already saved citations are visually differentiated with a distinct background color, giving users a quick visual cue indicating their ”as-read” status. This design enhances the user experience by efficiently tracking and managing saved citations, promoting a seamless transition between exploring new citations and revisiting previously saved content. Figures 3 b and c visually outline the process, illustrating how users can save inline citations and subsequently identify their saved status while exploring printed papers using the AR view.

#### 4.3.3. [DG3] Real-time Synchronization of Reading Notes

A cornerstone of AR-PaperSync’s functionality is the seamless synchronization of users’ reading notes, fostering a cohesive research experience across mobile, desktop, and printed paper formats. The system ensures real-time continuity, allowing users to take notes using the desktop application, with these notes dynamically synchronized across all platforms.

In the desktop application, users can meticulously annotate digital documents, highlighting essential sections and adding insightful comments to enhance their understanding. These reading notes are stored in the cloud service’s dedicated notes table, associating each note with the user, paper, and selected text. This robust backend infrastructure enables secure storage and efficient retrieval of users’ reading notes.

The real-time synchronization extends to the mobile AR environment, where users can witness their reading notes seamlessly appearing on printed papers in real time. Leveraging advanced OCR capabilities, the system extracts text from the printed paper, matching it with the highlighted text from the desktop application. When a match is found, the system dynamically prepares clickable highlighted text on the AR view, precisely overlaying the selected text on the physical document. Users can effortlessly click on these highlighted sections to reveal the associated comments from their desktop application.

This innovative approach ensures that users’ interactions with academic content, whether on a digital client or printed paper, remain synchronized and accessible in real time. Figure 5 illustrates the workflow of image processing to AR mobile view, and Figure 4 showcases how reading notes created on the desktop application seamlessly translate into highlighted sections on printed papers via the AR view, providing users with an integrated and fluid research experience.

### 4.4. Challenges and Solutions

During the development of AR-PaperSync, several intricate challenges arose that demanded innovative solutions for effective resolution. Capturing expansive images on mobile devices while maintaining optimal OCR accuracy required meticulous algorithmic optimizations, ensuring high-quality image processing without compromising text extraction fidelity. Additionally, translating inline citations and highlighted text from printed papers into an intuitive augmented reality format posed formidable challenges, necessitating the utilization of advanced AR technologies and user-centric design principles to facilitate seamless interaction. Enabling real-time data synchronization across disparate platforms accentuated inherent complexities, compelling the implementation of robust data synchronization protocols and cloud infrastructure capabilities to maintain data coherence and consistency. Furthermore, optimizing backend processing efficiency while enhancing rendering performance on the Mobile AR interface entailed strategic infrastructure scaling, algorithmic refinements, and performance tuning to deliver a responsive and immersive user experience across backend and frontend environments.

## 5. User Study

We present a comprehensive user study of AR-PaperSync, a system that innovatively combines augmented reality (AR) with traditional paper-based reading of scientific documents. This study is designed to critically assess the impact of AR-PaperSync on enhancing the academic reading experience, emphasizing aspects such as user interaction, time consumption, accuracy, usability, and cognitive load.

### 5.1. Research Hypotheses

Through a formal face-to-face usability study, we seek to answer the following pivotal research questions using the Goal Question Metric (GQM) [30] framework to define the goals of our study. This section outlines the research goals and corresponding hypotheses to guide our assessment of AR-PaperSync in enhancing the academic reading experience.

**Goal 1**: Assess how AR-PaperSync influences users’ efficiency and accuracy in accessing information compared to conventional methods.

Hypothesis 1: There is a significant improvement in efficiency and accuracy when using AR-PaperSync compared to conventional methods for accessing information in scientific printed papers.

**Goal 2**: Investigate the impact of AR-PaperSync on the usability and cognitive load during academic reading activities.

Hypothesis 2: AR-PaperSync reduces cognitive load and enhances the usability of academic reading activities compared to traditional methods.

**Goal 3**: Examine the usefulness of AR-PaperSync features in enhancing the reading experience of printed scientific papers.

Hypothesis 3: AR-PaperSync’s features significantly improve the reading experience of printed scientific papers compared to reading without AR-PaperSync.

### 5.2. Participants

The study involved 28 participants, 17 males and 11 females, with an average age of 26 *±* 4 years. The participant group comprised 15 PhD and 13 master’s students from the university. Participants, all of whom were enrolled in CSCI 488/688 courses, received one credit point towards their final score in these courses for their participation in the study. Among the participants, 18 reported regularly reading physical and digital scientific papers. In contrast, 3 participants primarily used printed materials, and 7 preferred exclusively with digital papers. None of the participants had prior experience with reading papers using an AR reading tool. This study was conducted as human subjects research and received approval from the Institutional Review Board (IRB). Each study session was conducted face-to-face, lasting between 20 and 30 minutes, and took place in the Department of Computer Science, specifically in the room ’QBB A5’.

### 5.3. Experimental Design

In our study, twenty-eight participants were evenly divided into two groups to assess the effectiveness of AR-PaperSync compared to traditional reading methods. Group 1 utilized AR-PaperSync, an innovative system integrating augmented reality on mobile devices to enhance interaction with physical documents and desktop applications. Group 2, employing the Baseline method, relied on traditional printed scientific papers and standard digital devices like computers or mobiles. Both groups were tasked with three specific activities during their reading sessions: finding inline citations, saving important citations, and taking reading notes. The experiment was conducted in a controlled environment using standardized equipment, including a MacBook Pro 15 inches, an Android phone equipped with the AR-PaperSync app, and a carefully selected short paper from the CHI EA archives, chosen for its readability and suitability for the participant’s academic levels.

### 5.4. Procedure

The experimental procedure was meticulously planned to ensure a fair and effective comparison between AR-PaperSync and the Baseline method. Upon arrival, participants provided formal consent and were given a brief orientation to familiarize them with the study’s objectives and the involved technology. Group 1 participants were introduced to AR-PaperSync and received a tutorial on utilizing its AR features on mobile devices. Conversely, Group 2 participants were briefed on the Baseline method, which entailed traditional paper-based reading supported by computers or mobile devices.

Participants were then instructed to read the provided CHI EA printed paper comfortably. Then, they engaged in the reading tasks, which involved finding inline citations, saving necessary citations, and taking reading notes on the paper. These tasks mirror typical academic reading scenarios, ensuring participant interactions were as natural and authentic as possible. For finding inline citations, Group 1 used the AR-PaperSync app to locate and read referenced papers, while Group 2 searched for referenced paper titles on the internet. To save necessary citations, Group 1 could save and view them in AR via the AR-PaperSync app, whereas Group 2 marked them on printed paper or saved them on their computer. For taking reading notes, Group 1 utilized the AR-PaperSync desktop tool for note-taking and the mobile app for viewing notes alongside the printed paper. Group 2 participants took notes directly on the printed paper or a computer.

Upon completing the three tasks, participants were asked to complete the System Usability Scale (SUS) and NASA Task Load Index (NASA-TLX) questionnaires for usability evaluation. Participants who used AR-PaperSync also provided qualitative feedback focusing on their perceptions, usability experiences, and potential improvements for the system. This feedback was gathered through the following interview questions:

1. How are the AR-PaperSync features useful to support the reading experience on printed paper?
2. Are there any AR-PaperSync features you will use in the future if the application is available?
3. Is there anything you want to improve or add to the AR-PaperSync?

### 5.5. Data Collection

Our study’s data collection and analysis process was meticulously designed to be comprehensive and systematic, capturing both quantitative and qualitative aspects of user interactions with AR-PaperSync and the Baseline method.

#### Quantitative Data Collection

In the quantitative phase of our study, meticulous attention was given to recording each participant’s time to complete specific tasks, including finding inline citations, saving necessary citations, and taking reading notes. This approach allowed for a precise measurement of task completion efficiency. Additionally, we paid close attention to the accuracy with which these tasks were executed, particularly in correctly identifying and saving citations and notes. Upon completing these tasks, participants were asked to fill out the System Usability Scale (SUS) and the NASA Task Load Index (NASA-TLX) questionnaires. These instruments provided valuable quantitative measures of usability and cognitive load, respectively. The data collected from these sources were integral to evaluating the functional effectiveness of AR-PaperSync compared to the Baseline method.

#### Qualitative Data Collection

The qualitative dimension of our study concentrated on acquiring more profound insights into the user experience and perceptions of AR-PaperSync. This was accomplished through a series of interviews, where participants responded to three critical questions outlined in section 5.3. These questions were carefully crafted to probe the usability and practicality of AR-PaperSync’s features. The feedback from these interviews was instrumental in understanding aspects of the user experience that extended beyond the quantitative data. In addition to these interviews, observational notes were taken during the tasks to document user behaviour, challenges faced, and overall engagement with the system.

## 6. Results

We evaluate the AR-PaperSync system by thoroughly examining quantitative and qualitative outcomes and comparing them to a baseline. These assessments aim to empirically confirm the system’s impact on the user experience during academic reading tasks.

### 6.1. Quantitative Evaluation

*Search Time.* Our evaluation began with a thorough examination of search time, revealing a significant reduction in task completion duration when AR-PaperSync was tested. Statistical analyses were employed to validate the system’s clear superiority over the baseline. On average, participants demonstrated remarkable efficiency, completing tasks a stunning 90% faster with AR-PaperSync compared to the baseline system. Notably, the analysis uncovered substantial improvements in search time, particularly in inline citations. While the baseline required an average of 30 seconds for this task, AR-PaperSync accomplished it in a mere 9 seconds. Similarly, the time taken for reading notes and saving citations was reduced by 70% and 40%, respectively, when AR-PaperSync was compared to the baseline. Paired-sample t-tests provided robust evidence of AR-PaperSync’s efficiency, consistently demonstrating significantly reduced search times compared to the baseline across all tasks: inline citations *(t-statistic = -6.21, p ¡ 0.001)*, reading notes *(t-statistic = -5.78, p ¡ 0.001)*, and saving citations *(t-statistic = -3.85, p ¡ 0.001)*. For a comprehensive breakdown of search time across these three tasks, please refer to figure 6.

**Figure 6:**
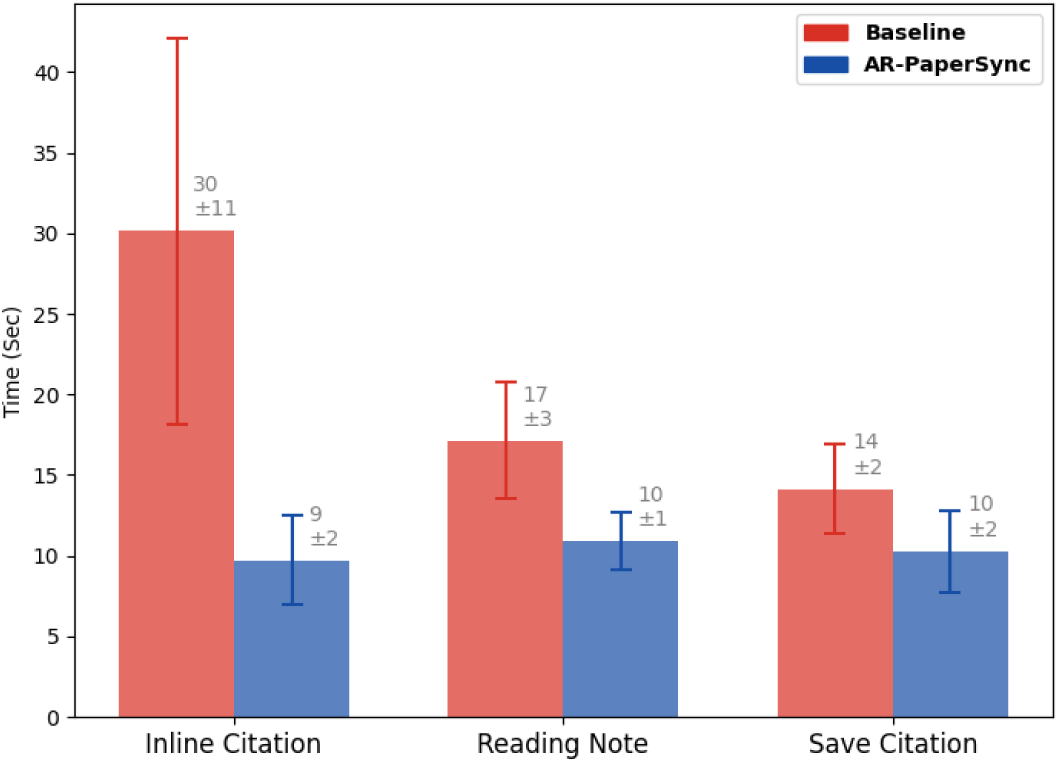
Comparative analysis of search time across inline citations, reading notes, and saving citations with AR-PaperSync and the baseline system.

#### Accuracy

Accuracy in task completion was significantly improved with AR-PaperSync. We used paired-sample t-tests to establish that the system outperformed the baseline in saving inline citations and note-taking activities. With AR-PaperSync, the average accuracy for reading notes and saving citations reached 91% and 94%, respectively, representing a 10% improvement over the baseline. Notably, inline citations saw the most substantial accuracy enhancement, with AR-PaperSync achieving near-perfect performance at 97%, while the baseline lagged behind at 80%. For a detailed performance comparison between AR-PaperSync and the baseline, refer to figure 7. Paired-sample t-tests revealed that AR-PaperSync exhibited higher accuracy than the baseline in inline citations *(t-statistic = 4.26, p ¡ 0.001)*, saving citations *(t-statistic = 2.13, p ¡ 0.05)*, and reading notes *(t-statistic = 1.47, p ¡ 0.1)*.

**Figure 7:**
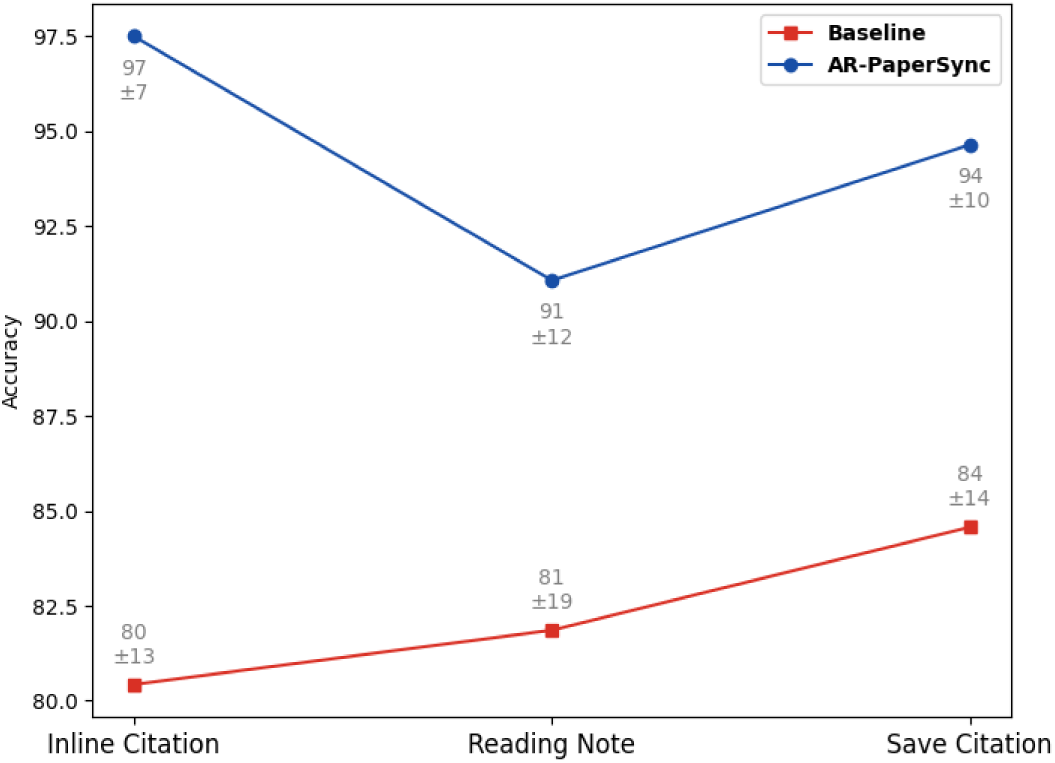
Performance comparison between AR-PaperSync and baseline in various task categories, including inline citations, saving citations, and reading notes

#### Cognitive Load

Our examination of cognitive load dimensions, assessed through NASA-TLX measures, unveiled compelling outcomes. AR-PaperSync consistently showcased its remarkable capacity to reduce mental workload across various dimensions significantly. Participants experienced substantial alleviation in mental, physical, and temporal demands, accompanied by heightened perceived performance, focused effort, and reduced frustration levels when engaging with AR-PaperSync. The average NASA-TLX score for AR-PaperSync stands at 18, underscoring a comprehensive enhancement in cognitive load metrics compared to the baseline, which was noted at 46. Further reinforcing this, a paired-sample t-test robustly confirms that the overall cognitive load is markedly lower for AR-PaperSync compared to the baseline *(t-statistic = - 16.51, p ¡ 0.001)*. Individual TLX subscales further highlight AR-PaperSync’s effectiveness in diminishing mental *(t-statistic = -9.85, p ¡ 0.001)*, performance *(t-statistic = -8.60, p ¡ 0.001)*, and frustration *(t-statistic = -8.40, p ¡ 0.001)*. Detailed NASA-TLX subscale scores are visually presented in Figure 8 for comprehensive insight.

**Figure 8:**
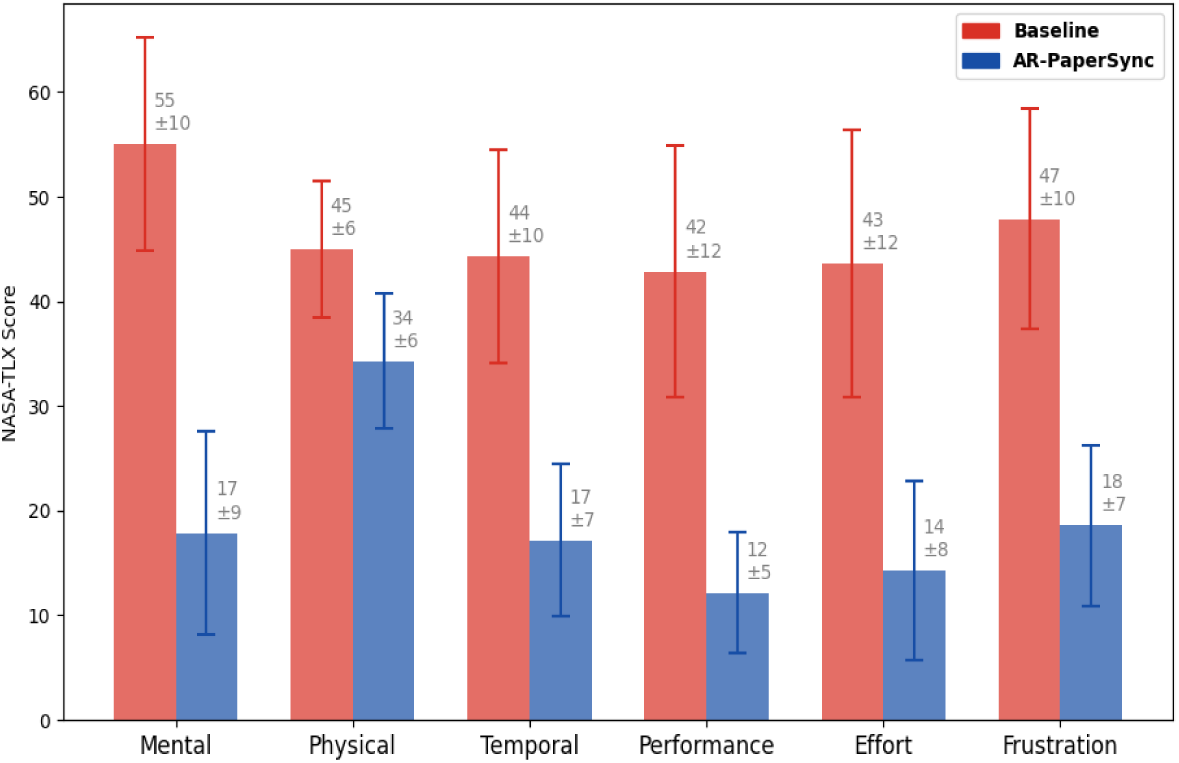
NASA-TLX subscale scores for AR-PaperSync and baseline comparisons.

#### System Usability

Our evaluation extended to assessing the system’s usability, as quantified through the System Usability Scale (SUS). The results painted a striking picture of user satisfaction and ease of use when engaging with AR-PaperSync. Impressively, AR-PaperSync garnered substantially higher SUS scores compared to the baseline. These elevated scores indicate a system that users found remarkably intuitive and efficient. The average SUS score for AR-PaperSync reached an impressive high of 78, more than 59% of the base-line score, underscoring the system’s user-friendly nature and aligning with the positive findings related to task performance. The paired-sample t-test results further confirm the significant difference, with AR-PaperSync surpassing the baseline in terms of system usability *(t-statistic = 10.21, p ¡ 0.001)*. A visual representation of the SUS score comparison is thoughtfully presented in Figure 9.

**Figure 9:**
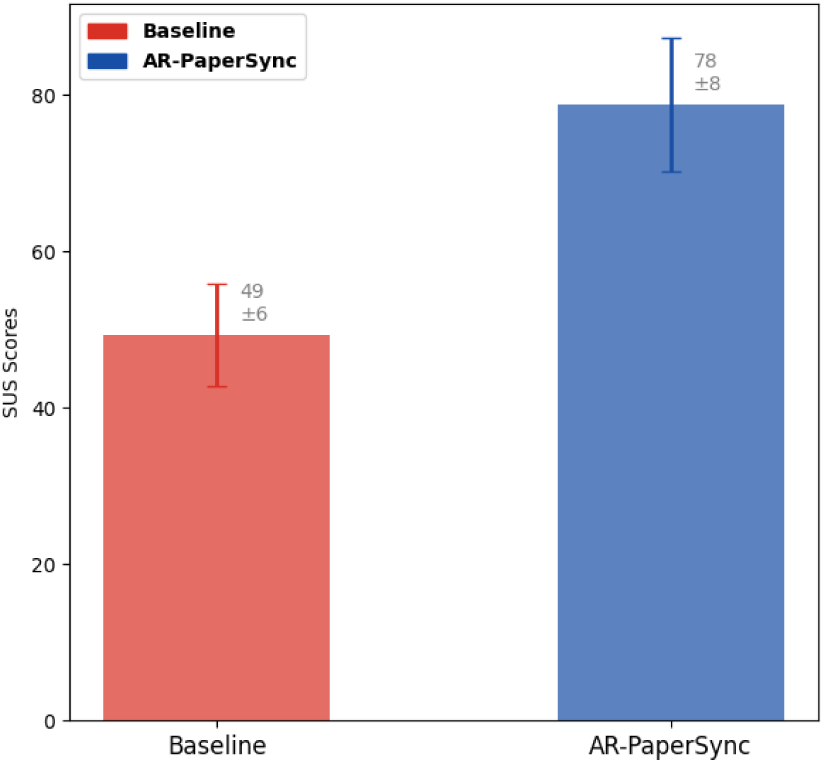
Comparison of System Usability Scale (SUS) Scores for AR-PaperSync and Baseline.

### 6.2. Qualitative Evaluation

#### Usefulness and Impact of AR-PaperSync Features

The feedback from participants demonstrates their appreciation of how AR-PaperSync has enriched their reading experience, offering significant practical advantages. The positive reception of AR-PaperSync’s features is striking, with 89% of respondents affirming its usefulness. Key terms that frequently emerged in participants’ comments about AR-PaperSync’s benefits include *”useful”*, *”good”*, *”greatly enhanced”*, *”easier”*, *”excellent”*, *”fast”*, and *”surprised”*. Specifically, Participant P9 effectively summarized the core benefit: *”Saves significant time and effort in locating papers”*. P1 highlighted its practical utility: *”Extremely useful for marking and annotating papers, offering time savings over traditional methods”*.

Participants P14 and P13 expressed a sense of wonder, noting, *”The experience was delightful, especially being surprised by the appearance of citations on printed paper”*, and *”It seamlessly integrates my online annotations for quick access via AR”*. The interaction with AR-PaperSync is likened to using a dynamic app instead of paper. *”Interacting with citations in AR was akin to engaging with a physical paper”*, and *”It substantially enhances digital engagement with printed materials”*, observed P2 and P5, indicating a blurring of the lines between physical and digital paper. The ease of navigating citations was also a focal point. P8 and P3 noted the efficiency advantage: *”Allows rapid checking of citations”* and *”Significantly speeds up the process of reading physical papers”*. Meanwhile, P6 praised the visual features: *”The system operates swiftly, making citations visible and easy to save”*. This collective feedback suggests that AR-PaperSync effectively tackles some of the most labor-intensive aspects of academic research, particularly in managing extensive bibliographies and notes.

#### Future Adoption and Use Intentions

User feedback regarding AR-PaperSync distinctly indicates a strong inclination toward future adoption and continued utilization of the system’s features. Remarkably, a substantial 93% of respondents affirmed their intent to utilize at least one feature offered by this tool in the future. Among these respondents, 74% expressed a keen interest in employing inline citations, 52% were inclined towards using reading notes, and 42% considered saving citation features appealing and practical. Notably, the sentiment was overwhelmingly positive, with P1 stating, *”I like inline citations and add comments option”*. This sentiment reverberated throughout several participants’ responses, with one participant outlining a specific use case: *”I find inline citation feature helpful during literature reviews when I need to refer back to the reference paper”*. Even users who prefer digital media expressed openness to incorporating physical media into their workflow, thanks to AR-PaperSync. P3 commented, *”I typically use digital media, but I can see myself embracing physical media with this application”*. This highlights the system’s potential to bridge the gap between digital and physical reading preferences. Furthermore, P5, P7, P12, and P14 emphasized the practicality and usefulness of AR-PaperSync features, such as marking read citations, adding comments, saving citations, and quickly accessing in-text citations in printed materials. These features were perceived as valuable for both current and future research endeavors. Overall, the user responses suggest a high likelihood of future adoption and continued use of AR-PaperSync, indicating that it can significantly impact and enhance reading and research practices.

#### Enhancements and Feature Requests

Collecting feedback on desired additional functionalities, P9 suggested *”Making open-source cited paper available with one click”*, indicating a need for streamlined access to resources. P10 envisioned more comprehensive integration: *”I wish to read cited papers remotely after saving, without having them with me”*, hoping for a feature allowing access to saved citations and notes beyond the immediate physical space. While the feedback was overwhelmingly positive, participants suggested improvements, such as increasing the app’s speed and introducing different functionalities.

## 7. Discussion

We discuss the implications of the AR-PaperSync study results, closely examining how the system aligns with our research hypotheses and the insights gathered from user feedback.

### 7.1. Efficiency and Accuracy in Accessing Information

The quantitative data from our study strongly support hypothesis 1, demonstrating that AR-PaperSync significantly enhances both efficiency and accuracy in accessing information. Participants using AR-PaperSync completed tasks 90% faster on average than the baseline system, reducing the time for finding inline citations from 30 seconds to just 9 seconds. This dramatic increase in efficiency is particularly crucial in academic research, where time is often scarce. Moreover, AR-PaperSync users achieved a 97% accuracy rate in identifying inline citations, significantly higher than the baseline’s 80%. The improvement in accuracy is vital, as it ensures that researchers can rely on AR-PaperSync for precise information retrieval. With AR-PaperSync, the average accuracy for reading notes and saving citations reached 91% and 94%, respectively, highlighting the system’s capability to make the academic reading process more efficient and precise.

### 7.2. Reduction in Cognitive Load and Enhanced Usability

Our study’s results from the NASA-TLX and SUS assessments robustly confirm hypothesis 2. Participants reported significantly lower cognitive load scores, with an average NASA-TLX score of 18 compared to the baseline’s 46. This reduction in cognitive load can be attributed to AR-PaperSync’s intuitive design and user-friendly interface, which streamline the traditionally complex process of academic reading. The high System Usability Scale (SUS) score of 78, significantly higher than the baseline, indicates that users found AR-PaperSync intuitive and straightforward. This ease of use is a critical factor in adopting new technologies in research settings, as it can significantly enhance the overall efficiency and productivity of academic work.

### 7.3. Improvement in Reading Experience

The qualitative feedback collected aligns closely with hypothesis 3, affirming that AR-PaperSync substantially improves the reading experience of printed scientific papers. Participants appreciated the system’s practical advantages, with terms like *”useful”*, *”efficient”*, *”easier”*, and *”greatly enhanced”* frequently mentioned. For example, Participant P9 noted the significant time and effort savings in locating papers, and P1 commended the system for marking and annotating papers more efficiently than traditional methods. The delight and surprise expressed by Participants P14 and P13 in seeing how citations appeared on printed paper through AR highlight the novel experience AR-PaperSync offers. The system’s ability to blend digital engagement with physical materials, as observed by P2 and P5, and the ease of navigating citations (P8 and P3) and quickly accessing in-text citations in the printed paper (P6) are testament to AR-PaperSync’s effectiveness in tackling the labor-intensive aspects of academic research. These user experiences underscore AR-PaperSync’s capacity to enrich the traditional educational reading process by introducing seamless and dynamic AR interactions, making the process more engaging and efficient.

In summary, AR-PaperSync demonstrates significant potential in transforming academic research, making it more efficient, less cognitively taxing, and more engaging. The system’s design effectively addresses researchers’ real-world challenges, offering a novel solution combining the best physical and digital worlds in academic reading and research.

## 8. Conclusion and Future Work

### 8.1. Conclusion

AR-PaperSync, an innovative augmented reality system, significantly advances the integration of digital technology with traditional academic reading practices. Through its intuitive design and functionality, AR-PaperSync has demonstrated its efficacy in streamlining the management of citations and notes in printed scientific papers. Our user study results indicate that AR-PaperSync enhances efficiency and accuracy in accessing information and significantly reduces cognitive load, thereby transforming the traditional academic reading experience into a more engaging and efficient process. The system’s ability to seamlessly blend printed papers’ physical tangibility with the dynamic interactivity of digital tools represents a notable contribution to educational technology and academic research tools.

### 8.2. Future Work

In future iterations of AR-PaperSync, we plan to harness the capabilities of Machine Learning (ML) and deep learning to enhance the system’s functionality and user experience significantly [31, 32, 33, 34, 35]. We will integrate a feature to access all inline citations of a paper with a single click on AR view addressing participant P9’s feedback. In alignment with P10’s vision, we will develop realtime synchronization of saved citations from printed paper to desktop, enabling remote access from any environment. To improve responsiveness, ML-driven optimization techniques will be employed to boost the application’s speed. We also aim to expand our system with advanced NLP and deep learning techniques for deeper information extraction and automation, offering sophisticated content analysis and contextual insights. This strategic integration of ML and deep learning technologies is poised to transform AR-PaperSync into a more adaptive, efficient, and powerful tool, advancing its impact on academic research and educational technology.

